# High-Throughput Identification of Genetic Variation Impact on pre-mRNA Splicing Efficiency

**DOI:** 10.1101/191122

**Authors:** Scott I. Adamson, Lijun Zhan, Brenton R. Graveley

## Abstract

Understanding the functional impact of genomic variants is a major goal of modern genetics and personalized medicine. Although many synonymous and non-coding variants act through altering the efficiency of pre-mRNA splicing, it is difficult to predict how these variants impact pre-mRNA splicing. Here, we describe a massively parallel approach we used to test the impact of 2,059 human genetic variants spanning 110 alternative exons on pre-mRNA splicing. This method yields data that reinforces known mechanisms of pre-mRNA splicing, can rapidly identify genomic variants that impact pre-mRNA splicing, and will be useful for increasing our understanding of genome function.

## Background

One of the main goals of personalized medicine is to understand how genetic variations between individuals impact health. Genetic variants can impact health in a number of different ways, one such way is through altering pre-mRNA splicing efficiency. Alternative splicing is a process that is important for regulatory function and a primary source of proteome diversity in humans [1]. Perturbations in splicing have also been shown to contribute to a number of different diseases [2,3]. These splicing changes can manifest themselves through interrupting well-known interactions between the spliceosome and splicing elements including the 3’ and 5’ splice sites, pyrimidine tract, or branchpoint sequences. However splicing can also be perturbed by disrupting other sequences known to modulate splicing. Exonic splicing enhancers and silencers (ESEs and ESSs), as well as intronic splicing enhancers and silencers (ISEs and ISSs) are examples of splicing regulatory elements that can be perturbed and result in different splicing outcomes. Modulation of these splicing regulatory elements have been shown to be disease associated (for a review see [4]). Thus understanding how both intronic and exonic variants impact splicing not only provides insights into the mechanisms of splicing, but also is important to understand the basis of certain genetic diseases.

Identifying variants that impact splicing regulatory elements and their splicing consequences are difficult to detect using conventional poly(A)+ RNA-seq alone because the variants are often spliced out of the mature mRNA. There have been a number of different studies that have aimed to address this issue. One approach has been the pursuit of deciphering the “splicing code” using computational techniques such as deep learning [5–7]. While these studies have yielded useful knowledge about splicing and do have predictive power, experimental confirmation of the behavior of these variants has been limited and the predictions are not perfect. Other groups have pursued the use of random sequences to understand the splicing code, however it is hard to integrate datasets with contextual transcriptome information (i.e. CLIP) when studying the splicing behavior of random sequences [8]. A more recent study tested a number of exonic disease-associated variants in parallel using a mini-gene system [9]. The approach was to observe the allelic ratio of reference to variant in a plasmid pool, and compare with the ratios observed from splicing outcomes. This approach is useful for studying exonic variants, but is unable to test intronic variants. Here we present a method that address some of these shortcomings using a barcoding approach called <u>V</u>ariant <u>ex</u>on <u>seq</u>uencing (Vex-seq). Vex-seq is capable of testing many exonic and flanking intronic variants for the same exon simultaneously.

## Results

We set out to develop a high-throughput reporter system to determine the impact of genomic variants on pre-mRNA splicing. Our general approach is to generate a library of test exons flanked by two common constitutive exons (Figure 1). The library was introduced into tissue culture cells followed by RT-PCR and sequencing to determine the splicing frequency of the test exon. Importantly, the reporters also contain a barcode sequence that serves as an identifier of which exon was present in the reporter so that it is possible to associate the pre-mRNA of origin in cases where the test exon was skipped.

**Fig. 1.**
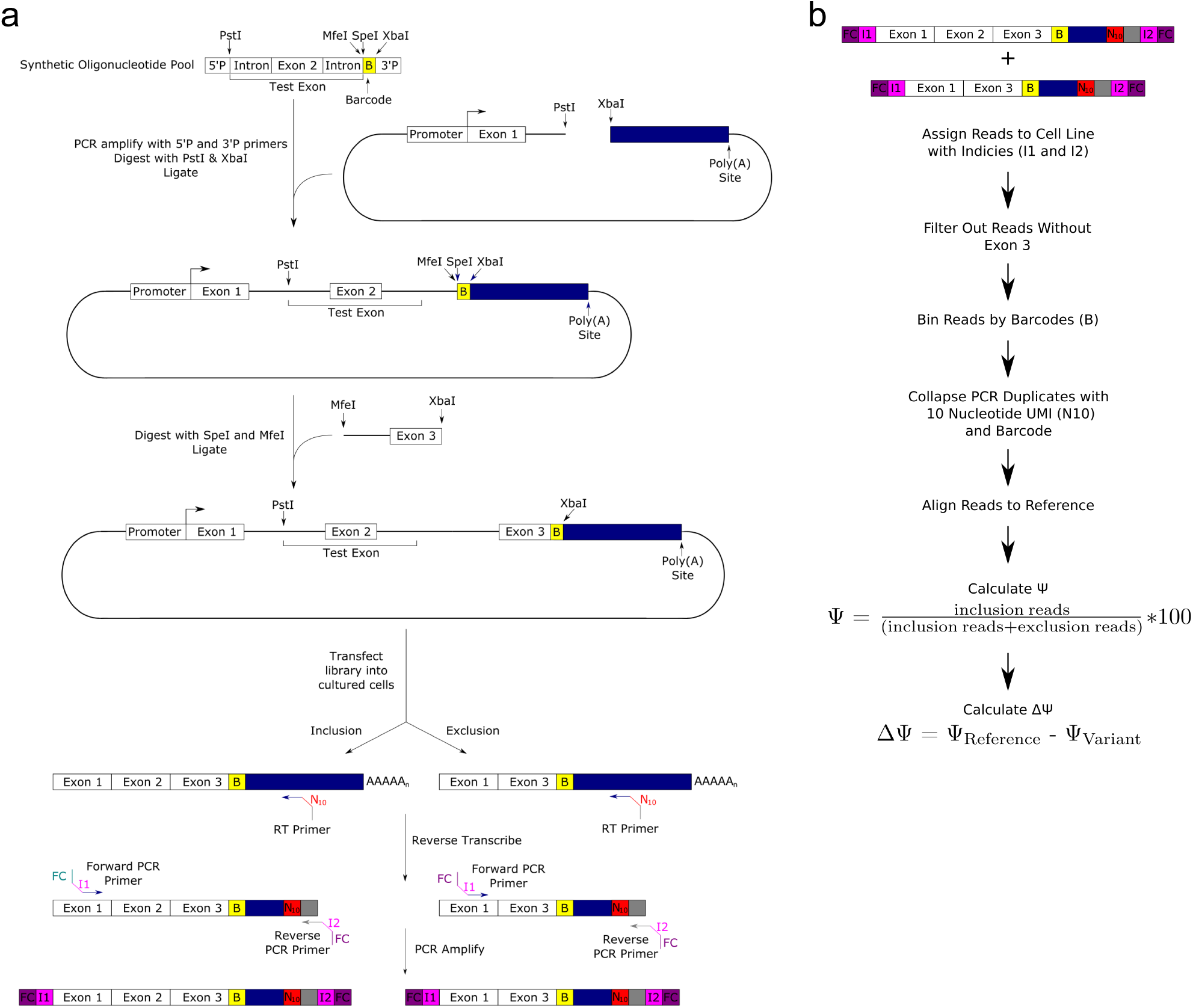
Assembly of test exon and experimental design. **a)** The test exon and flanking introns are subcloned into a reporter plasmid in a two step process, such that the barcode designating the sequence is near the end of the transcript. Once these plasmids are transfected into cultured cells, a transcript will be produced that may not contain the variant itself, but does contain the barcode (b) uniquely associated with the variant tested. A ten nucleotide UMI (N10) is attached during the reverse transcription step to collapse PCR duplicates downstream. Illumina flow cell binding sequences (FC) and indexes (I1 and I2) are attached via primers during PCR and the resulting DNA is sequenced on a MiSeq platform. **b)** Data analysis pipeline for splicing results.

We first designed a pool of 2,059 variants spanning 110 exons with reference, consensus splice site and mutated splice site control sequences for each exon. To insure reproducibility, each variant exon was associated with at least three unique eight nucleotide barcodes. Common primer sequences and restriction enzyme sites were also added for proper library construction. We included a minimum of 50 bases of the upstream intron, which should be adequate to include the majority of branchpoints [10], as well as the exon itself and at least 20 bases of downstream intron. This allowed for construction of test exons up to 97 nucleotides in length. Exons between the size of 68 and 97 nucleotides were randomly selected from Ensembl GRCh37.p13 annotations and variants from the ExAC database were intersected with the selected exons and their flanking intron sequences [11].

We amplified the oligonucleotide pool by PCR (Supp. Table 1). This product was then subcloned into a modified version of the splice reporter plasmid pcAT7-Glo1 in between the first intron and the 3’ UTR to generate a 1o library. Then restriction sites in between the barcode and the end of the test sequence were used to subclone in the second part of the second intron and third exon from the original plasmid (see Figure 1a). This results in a plasmid that encodes a transcript containing the first exon and part of the first intron of the globin gene, the test sequence, followed by the second intron and final exon of the reporter transcript, ending with the barcode and the 3’UTR. We refer to this final library pool as the 2o library.

In order to ensure that the variants are associated with the correct barcode, the 1o and 2o libraries were sequenced using paired end amplicon sequencing (Figure 2a). The results from sequencing the 1o library show that the majority of barcodes are correctly associated with the correct variant (Figure 2b). Barcodes excluded from the analysis due to having too few (less than 85%) of the correct variant reads associated with it only make up about 1.8% of the barcodes tested. Barcodes that were filtered out of the analysis also tended to have a lower read depth, suggesting that this may be related to the reason for their higher error rate (Figure 2c). We are also able to measure a misassignment rate of 4.59% using this plasmid sequencing technique. In order to ensure that the library contained a good diversity of sequences, we calculated a skew ratio between the 10th and 90th percentile of read depth for each barcode as done previously to check library diversity [12]. A skew ratio for the library established was calculated to be 5.5, which is considered adequate. We conclude that despite some misassignment, the plasmid pool is robust enough to be used to study variant changes in splicing.

**Fig. 2.**
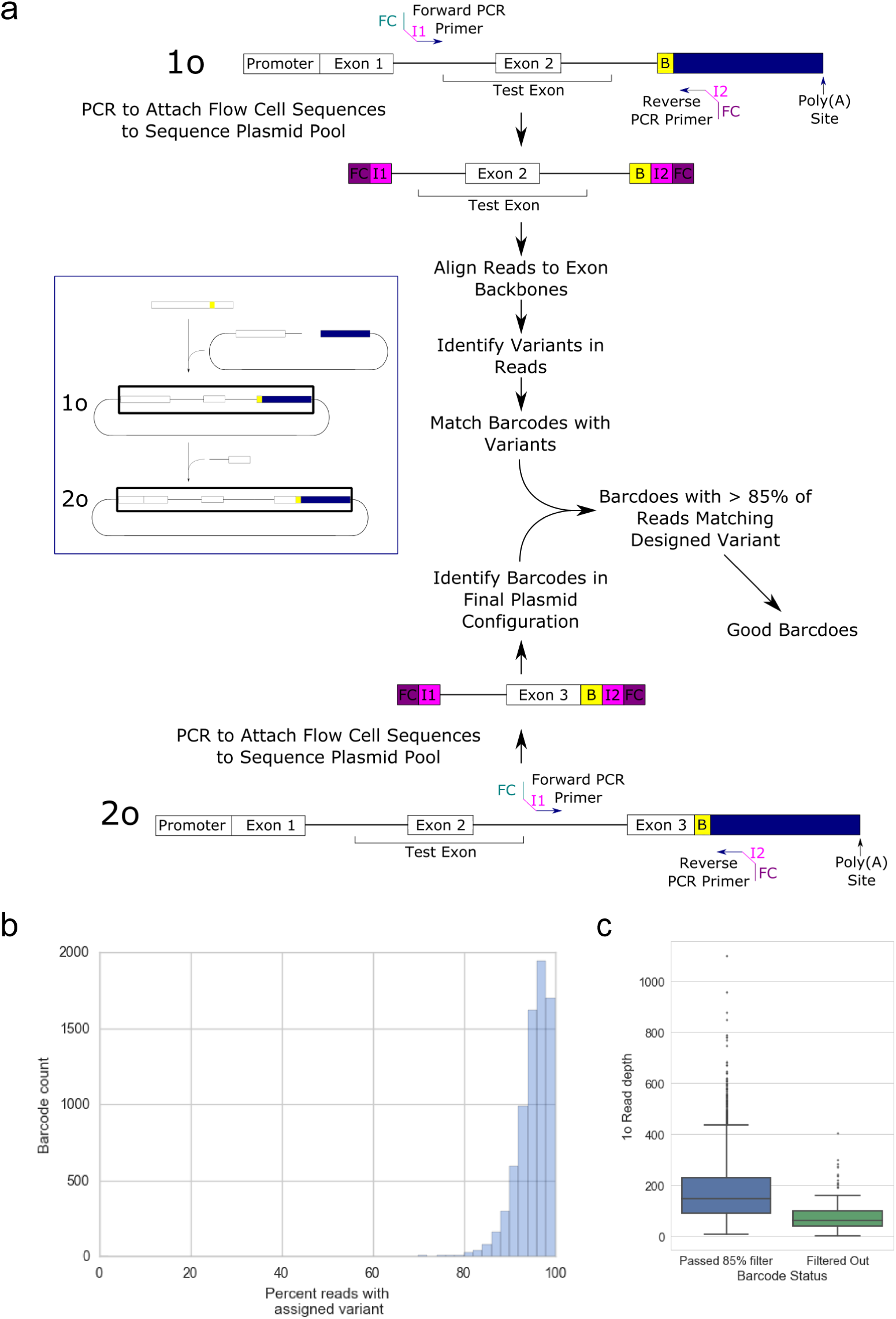
Quality control for plasmid integrity. **a)** Quality control pipeline for plasmid integrity. Amplicon sequencing of the 1o and 2o plasmid configurations are done through PCR to attach Illumina flow cell binding sequences (FC) and indexes (I1 and I2). Poor quality barcodes are then filtered out by identification of reads not containing variants and excluding barcodes with less than 85% of reads containing the correct variant. **b)** A histogram of the barcodes with the percentage of reads with correct variant identified. **c)** Box plots of 1o library read depth for barcodes included and excluded from further analysis.

The 2o library was then transfected into K562 and HepG2 cell lines in biological triplicate. cDNA was then synthesized from the RNA isolated from the cells using a mini-gene specific primer, a 10 nucleotide random sequence which serves as a unique molecular identifier (UMI) and an Illumina Read 2 sequencing primer. PCR amplified the cDNA to attach the other necessary sequences for Illumina paired end sequencing. The products were then sequenced on an Illumina MiSeq.

The data analysis pipeline uses custom python scripts to ensure that read 2 contains the third exon, the correct restriction site next to the barcode, and sorts the reads by barcode into bins. PCR replicate reads are collapsed into a single read using the UMI from the reverse transcription primer. The reads in each bin are then aligned using STAR to a reference specific to each variant [13]. Percent spliced in (PSI or Ψ) and change in PSI (ΔΨ) from the reference sequence are then calculated (Figure 1b). An amplicon based paired-end sequencing reads contain an unambiguous splicing outcome for each read, making Ψ and ΔΨ calculations straightforward for alignment outputs alone.

To assess how similar the barcodes associated with the same variant impacted the splicing behavior, we compared the Ψ value of the barcode replicates for each variant and reference exon and observed high correlations (Figure 3a). To ensure that these splicing values were robust to biological variation, we also examined the correlations of variants between 3 biological replicates for HepG2 and K562 cell lines (Figure 3b). This data shows similarly high correlation values between biological replicates of the same cell lines, showing robustness to biological variation. This data shows that the splicing data from Vex-seq is robust to both technical and biological variation.

**Fig. 3.**
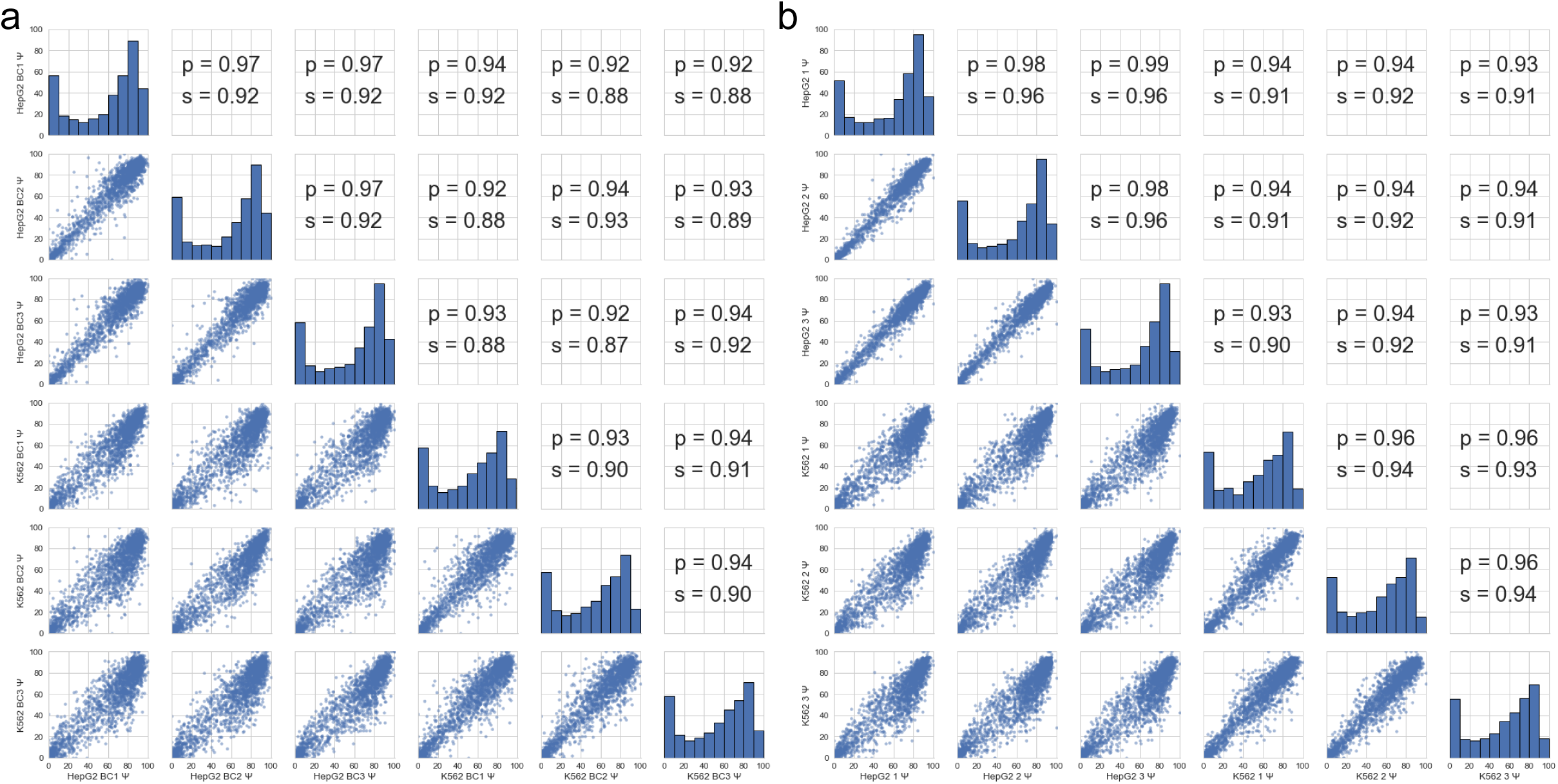
Behavior and reproducibility of of splicing outcomes. **a)** Spearman (s) and Pearson (p) correlations of barcode Ψ for the same variants. **b)** Spearman and Pearson correlations of biological replicate Ψ for each variant.

In order to ensure that splicing behavior reflects what is known about splicing, we examined the Ψ of the mutated and consensus splice site controls relative to reference and variant exons. For the mutated splice site controls, both splice sites were mutated such that the 3’ splice sites were changed from AG to TC and the 5’ splice sites were changed from GT to CA. For the consensus splice site controls, the variants contained a 20 nt pyrimidine tract, an AG at the 3’ splice site, and a consensus 5’ splice site of GTAAGT. The consensus and mutated splice site controls behave as expected with the mutated splice site controls displaying a low Ψ value and consensus splice sites having a higher Ψ value, while the variant and reference sequences are intermediate (Figure 4). These are consistent with expected splicing behaviors for these control sequences.

**Fig. 4.**
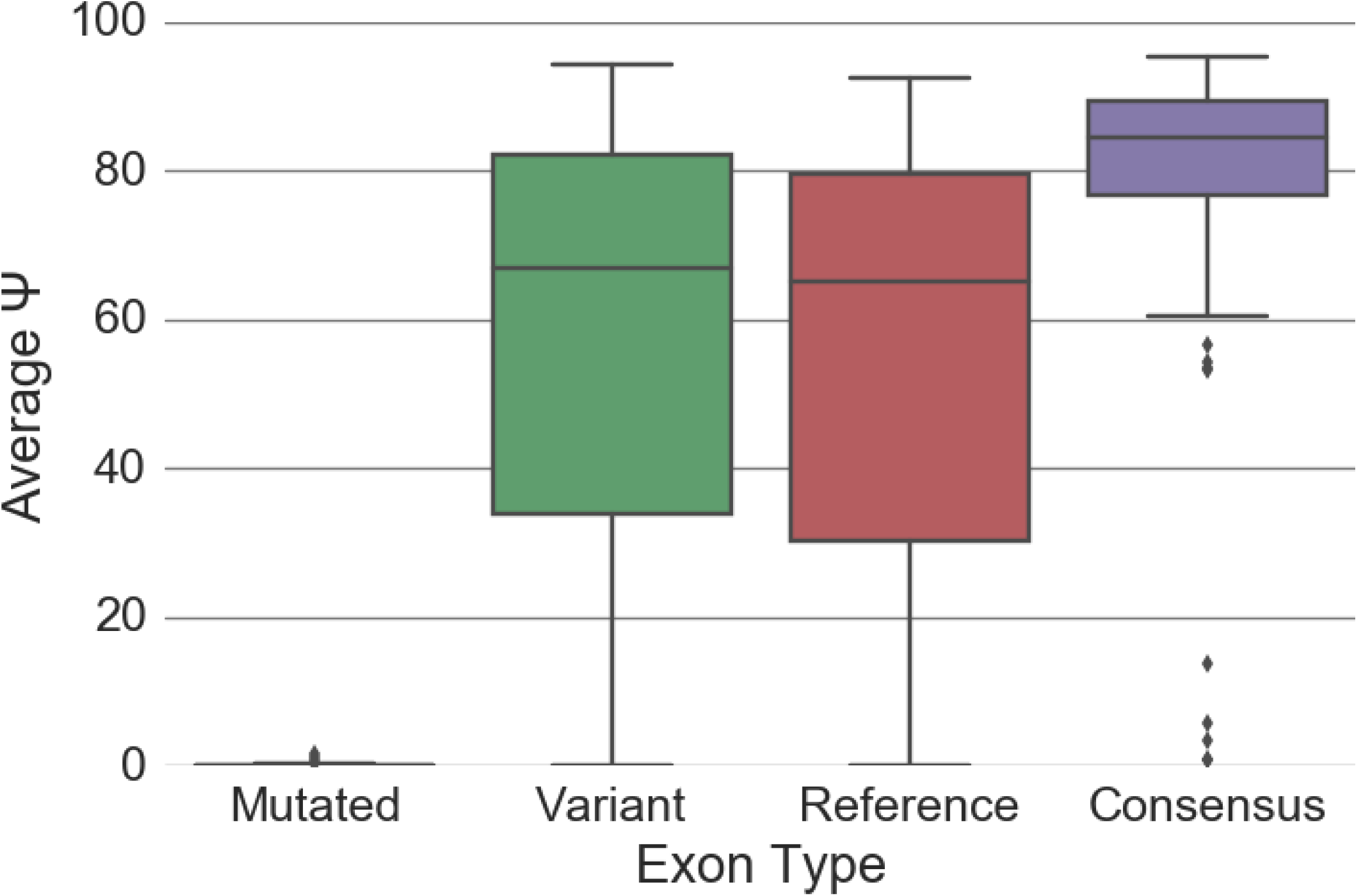
Box plots of mean Ψ for each type of control and test sequences.

Given the high correlation rates of Ψ values between the biological replicates of different cell lines, we sought to characterize this similarity further. Indeed, upon examining the correlation between the average Ψ value of each cell line, we observe a similar pattern (Figure 5a). Similarly, when looking at changes in splicing (ΔΨ) we see a similar, but noisier trend (Figure 5b). This suggests that variant induced changes in splicing studied in Vex-seq are generally cell type independent.

**Fig. 5.**
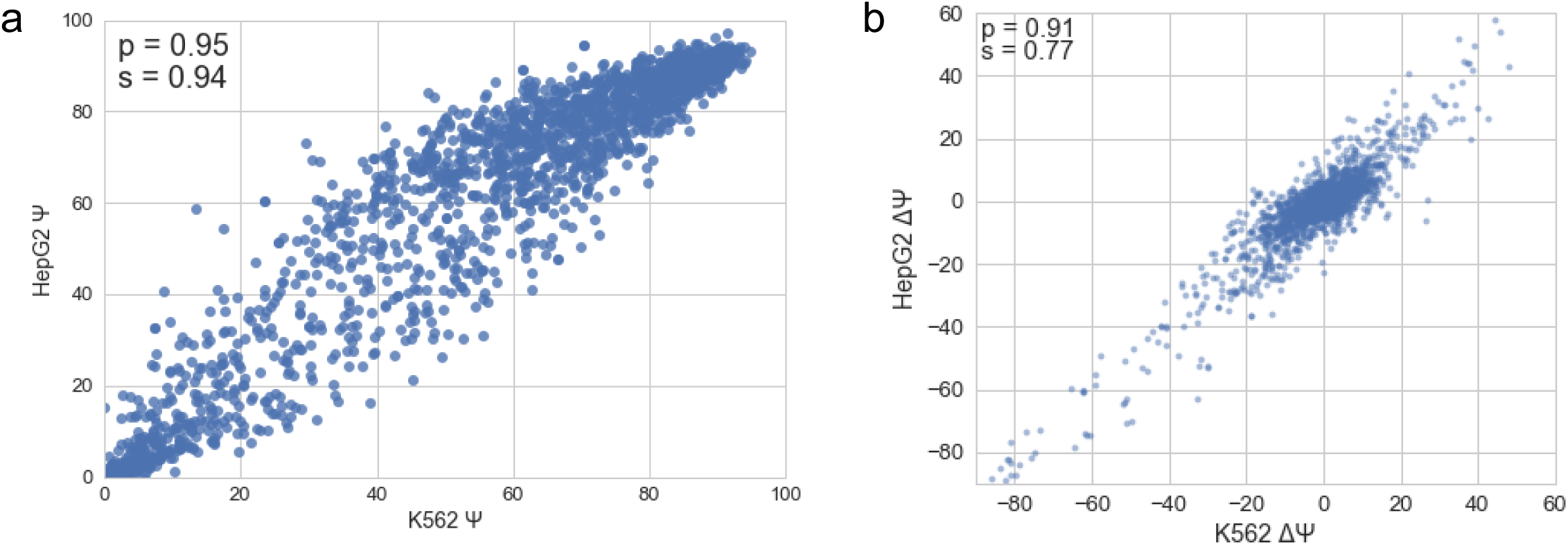
Similar behavior of splicing between K562 and HepG2 cell lines. **a)** Correlation between Ψ values for each variant between K562 and HepG2 cell lines. **b)** Correlation between ΔΨ values between K562 and HepG2

We next examined the changes in splicing efficiency (or ΔΨ) for each variant. Changes in splicing can be observed in many different positions relative to each of the splice sites, however perturbations in the 3’ and 5’ splice sites typically result in a dramatic reduction of ΔΨ (Figure 6). Many outliers can be observed changing ΔΨ negatively upstream of the 3’ splice site, which may correspond to changes in the pyrimidine tract or the branchpoint sequences. However variants in core splicing sequences alone do not account for the full diversity of splicing variation observed from this data. Evidence of potential ESS and ESE regulatory elements can be observed within the exon, as variants in the exon are capable of inducing strong ΔΨ changes in either directions.

**Fig. 6.**
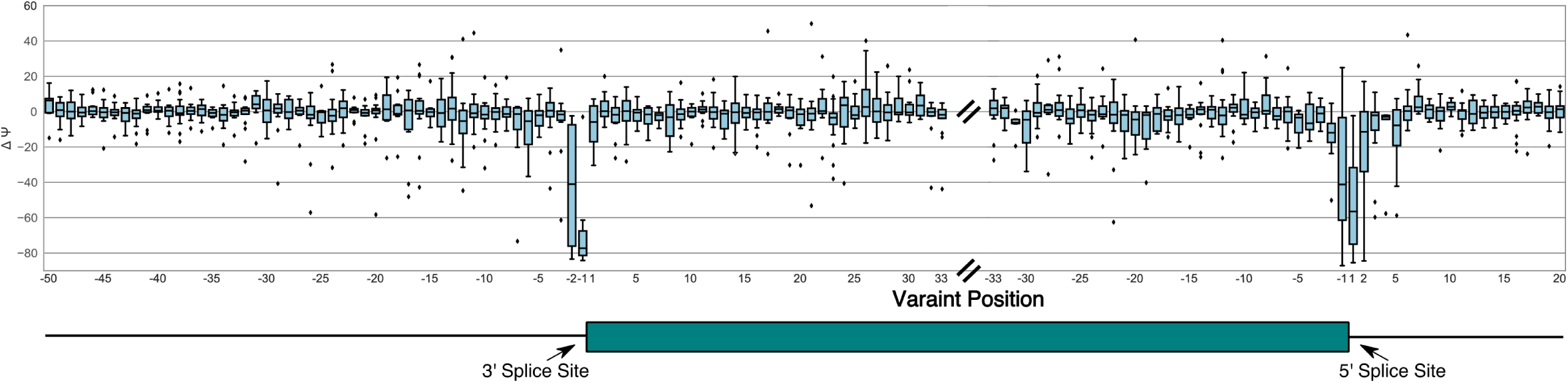
ΔΨ from both K562 and HepG2 plotted for all variants relative to the 3' and 5' splice sites.

To further characterize variants being studied and how they impact splicing we looked at other features including effect predictions. Using Variant Effect Predictor (VEP) [14] annotations we characterized the variants and their impact on ΔΨ (Figure 7). Annotations were selected based on the first reported annotation from VEP. The 5’ and 3’ splice site variants have the biggest negative impact on ΔΨ as expected. Intron, missense, synonymous, intron and splice region variants can also have a wide range of impacts on ΔΨ. This is consistent with previous findings about how missense and synonymous variants can change splicing inclusion levels [15]. It should also be noted that splice region variants alone do not account for many of the variants which changed splicing, consistent with the difficulty of predicting the impact of these variants based on impact annotations.

**Fig. 7.**
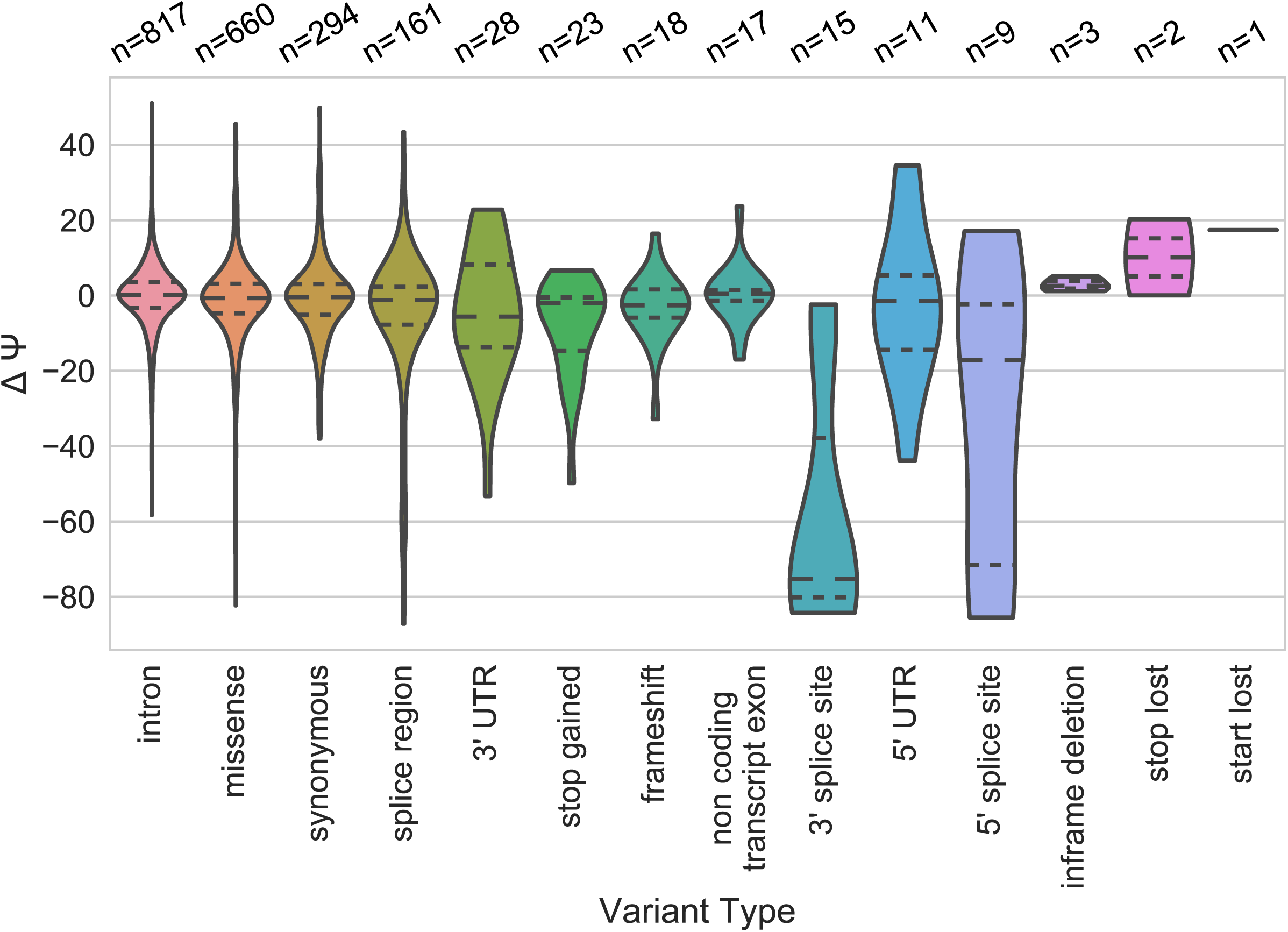
Variants classified by effect prediction and their impact on ΔΨ.

We were also interested in examining whether the variants that displayed the largest ΔΨ were more or less conserved than variants that had little impact on ΔΨ. 100-way vertebrate conservation scores from PhyloP were used to examine how variants with impacts strong or weak impacts on splicing were conserved [16]. We observed that there is significantly more conservation in variants which tend to have a high impact on splicing (|ΔΨ| ≧ 5) compared to variants with a low impact on splicing (|ΔΨ| < 5) (Figure 8a). Much of the conservation observed is likely due to protein coding constraints on sequences, which may add noise to this signal. To investigate whether this splicing-sensitive conservation is stronger in variants without protein changing potential, we examined the same trend in variants without protein coding constraints (intron, synonymous, UTR, and splice region variants), and we observed a more significant difference (Figure 8b). Additionally, when we focus on synonymous variants only, the effect is much clearer, even with a smaller sample size (Figure 8c). Intron variants seem to show the same trend of higher conservation with higher |ΔΨ|, however it is a milder effect (Figure 8d). This suggests that ESEs and ESSs modulated by these variants are more conserved, while intronic regulatory regions in the window we are testing are relatively more flexible. Perhaps this weaker conservation signal because ISSs and ISEs are not constrained by the context of the protein frame, and may be able to move around in linear space within the intron and still be effective in influencing splicing.

**Fig. 8.**
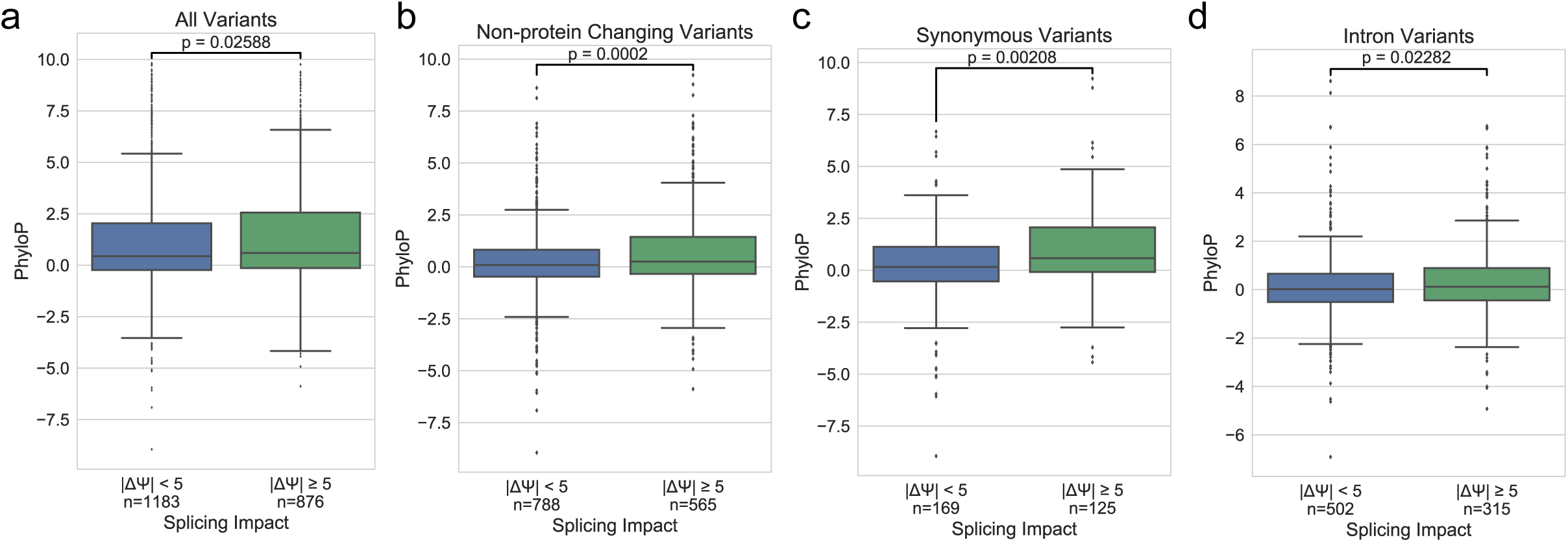
Conservation of variants with strong splicing impacts. **a)** Boxplots showing the relationship of PhyloP and magnitude of ΔΨ for all variants. **b)** Boxplots showing the relationship of PhyloP and magnitude of ΔΨ for variants without explicit protein coding annotation. **c)** Boxplots showing the relationship of PhyloP and magnitude of ΔΨ for synonymous variants. **d)** Boxplots showing the relationship of PhyloP and magnitude of *ΔΨ* for intron variants. P-values are calculated using Mann-Whitney-U test.

## Discussion and Conclusions

We have developed a new assay to assess how variants can impact pre-mRNA splicing efficiency called Vex-seq. This method builds upon previous high throughput splicing reporter assays. It utilizes a barcoding approach and designed sequences based on the transcriptome. This approach of using designed sequences allows for the possibility of not deeply sequencing the plasmid pool, because barcode variant associations are already known. This assay is able to test intronic variation which other recent methods have been unable to do [9]. Vex-seq is also able to account for the impacts that variants may have on transcription of reporter transcripts because of the barcoding approach. This assay could be applied to a number of different applications including fine mapping of GWAS variants that may be involved in splicing regulation, which has been shown to be linked to complex diseases [3]. Additionally this could be used to dissect the behavior of RNA binding proteins and their effect on splicing regulation, or even saturating mutagenesis of exons known to be important for diseases. Thus, Vex-seq has the potential to have an extremely high impact on our understanding of genome function and how non-coding sequence variants impact pre-mRNA splicing.

While Vex-seq offers certain advantages over current methods, there remain some obstacles with all of these splice reporter approaches [8,9]. First, these massively parallel splicing assays lack the context of the entire gene and chromatin state of the native genes. Second, these assays have limitations in terms of barcode design and synthesis length constraints and also may have cryptic splice sites formed in the context of the mini-gene. As oligonucleotide synthesis technologies improve, more context can be added to exons tested in this way. With more context, we expect Vex-seq to be more accurate at identifying variants that impact splicing. Third, these approaches also have some difficulty accounting for nonsense mediated decay, which may bias different variants compared to their impact in the context of the transcriptome.

Despite only examining 110 alternative exons in this study, we are able to obtain some biological insights from this data. The first is the similarity between the splicing behavior of K562 and HepG2 cell lines. 76.45% of variants agree in directionality of ΔΨ between cell lines, furthermore, when you restrict this analysis to only include variants that have a |ΔΨ|> 5 in HepG2, the agreement is even stronger (92.61%). This observation is consistent with the well correlated Ψ and ΔΨ values for variants (Figure 5). This may suggest that most variants tested in this context are acting upon splicing elements common across these cell lines. This observation may change when analyzing splicing changes in response to stimuli or in the context of a cells with more complex transcriptome regulation. Alternatively, this may suggest that regulatory factors important for cell type specific splicing are generally outside of the window that we are testing in Vex-seq. The predictive power of conserved intronic splicing regulatory elements on Ψ generally being within 100 nucleotides upstream and downstream may suggest that this is the case [17].

Data obtained from Vex-seq demonstrates the importance of variants on impacting pre-mRNA splicing efficiency. It shows that variant effect prediction, while useful for predicting protein changing variants, is insufficient to predict all splicing changes induced by variants. We also show that variants that tend to change splicing more are also generally more conserved than nucleotides that do not, particularly when the variants are otherwise not predicted to change protein products.

## Methods

### Plasmid alterations

pcAT7-Glo1 was provided by Kristen Lynch. To eliminate a splice acceptor site in the middle of intron 1, a deletion of the pyrimidine tract and splice acceptor sequence was deleted. This was done through digestion of the vector with AflII and PstI and PCR amplifying an insert using two primers (FWD 5’-AAACTCTTAAGCTAATACGACTCACTATAGG -3’) (REV 5’-GACTGAATGAGTCTGCAGAGGCAGAGAGAGTCAGTGG -3’). The insert digested with AflII and PstI and was ligated in the vector digested with the same enzymes resulting in the plasmid used for these studies.

### Assembly of Vex-seq plasmid

The oligo pool (Supp Table 1) was amplified with a common primer set (FWD 5’- GTAGCGTCTGTCCGTCTGCA -3’) (REV 5’- CTGTAGTAGTAGTTGTCTAG -3’) for 20 cycles, then digested with PstI and XbaI. These were subcloned into the modified pcAT7-Glo1 also using PstI and XbaI sites. The resulting plasmid pool, referred to as 1o was then digested with SpeI and MfeI. Exon 3 and intron 2 were PCR amplified from the original plasmid with primers (FWD 5’- GTGTGGAAGTCTCAGGATCG -3’) (REV 5’- AACGGGCCCTCTAGAGC -3’) and digested with MfeI and XbaI. The resulting product was subcloned into the digested 1o vector resulting in the final plasmid pool (2o).

### Transfection and Cell culture

HepG2 cells were grown to a density of 0.5 x 10^6^ cells per well and transfected with 1 μg of plasmid DNA using Lipofectamine 2000. Transfected HepG2 cells were then selected with 1 mg/mL zeocin for 8 days. K562 cells were grown to a density of 1.0 x 10^6^ cells per well and electroporated with 5 μg of plasmid DNA. Transfected K562 cells were then selected with 200 μg/mL of zeocin for 8 days. RNA from each cell line was isolated using Maxwell® 16 LEV simplyRNA Purification kits.

### Sequencing Preparation

Sequencing for the 1o library was constructed using a nested PCR reaction. The 1o library was amplified for 15 cycles using the following primers (FWD 5’- ACACTCTTTCCCTACACGACGCTCTTCCGATCTCCACTGACTCTCTCTGCCTC -3’) (REV 5’-GTGACTGGAGTTCAGACGTGTGCTCTTCCGATCTAGCGGGTTTAAACGGGCCCT -3’). The 2o library was amplified for 15 cycles using the following primers (FWD 5’- ACACTCTTTCCCTACACGACGCTCTTCCGATCTAGCAGCTACAATCCAGCTACCA -3’) (REV 5’- GTGACTGGAGTTCAGACGTGTGCTCTTCCGATCTAGCGGGTTTAAACGGGCCCT-3’). Each of these products were then amplified for 10 cycles using the following primers (FWD 5’- AATGATACGGCGACCACCGAGATCTACAC*-i5-INDEX-* ACACTCTTTCCCTACACGACGCTCTTCCGATCT -3’) (REV 5’- CAAGCAGAAGACGGCATACGAGAT*-i7-INDEX-* GTGACTGGAGTTCAGACGTGTGCTCTTCCGATCT -3’). The cDNA was synthesized from the K562 and HepG2 RNA using SuperScript(™) III reverse transcriptase and a gene specific primer (5’- GTGACTGGAGTTCAGACGTGTGCTCTTCCGATCTNNNNNNNNNNGCAACTAGAAGGCACAGTCGAGG -3’). The cDNA was then PCR amplified for 10 cycles using primers (FWD 5’- AATGATACGGCGACCACCGAGATCTACAC*-i5-INDEX* ACACTCTTTCCCTACACGACGCTCTTCCGATCTGGCAAGGTGAACGTGGATGAAG -3’) (REV 5’- CAAGCAGAAGACGGCATACGAGAT*-i7-INDEX-*GTGACTGGAGTTCAGACGTGTGCTCTTCCGATCT -3’). Resulting samples were multiplexed and sequenced on a MiSeq using a v2 300-cycle kit.

### Data Analysis and Interpretation

#### Plasmid Quality Control

Forward and reverse reads from plasmids were combined into a single read using FLASH [18]. 1o library reads were sorted into bins using the barcode and grouped by control exon backbone with separate bins for indels and control sequences. Reads were then aligned using Novoalign V3.02.13 (http://www.novocraft.com). Sam2tsv was then used to identify variants in each read and identify the barcode sequence [19]. Barcodes with 15% or more of reads not containing the correct variant were filtered out during splicing analysis using custom python scripts. Barcodes were identified 2o library reads using custom python scripts and barcodes without reads were filtered out of the analysis.

#### Splicing Alignments and Analysis

Reads were identified by barcode and sorted into bins for each variant. The duplicate reads in each bin were then collapsed into a single read by the UMI. Reads were then aligned to a variant specific reference using STAR version 2.5.2b [13]. The uniquely aligned annotated read junctions were identified and Ψ and ΔΨ were calculated. Reads which spanned unannotated splice junctions were discarded for calculating Ψ and ΔΨ. Ψ values for analysis, unless otherwise indicated were the mean of the K562 and HepG2 Ψ values. Mutated and Consensus splice site controls were removed for the correlation analyses corresponding to figures 3A, 3B and 5. Annotations for each variants was done using the Ensembl Variant Effect Predictor tool using assembly GRCh37.p13 and using the Ensembl transcript database [14]. The variants used in the analysis were selected based on the first annotation output by VEP. 100-way vertebrate PhyloP conservation scores were used to examine conservation [16].

## Declarations

## Acknowledgments

We thank Kristen Lynch for the pcAT7-Glo1 plasmid and members of the Graveley lab for discussions.

## Funding

This work was supported by a grant from the National Human Genome Research Institute (R21HG008799) and the John and Donna Krenicki Endowment Fund to BRG.

## Availability of data and materials

The datasets generated and/or analysed during the current study are available in the Short Read Archive repository, [PERSISTENT WEB LINK TO DATASETS].

## Competing interests

The authors declare that they have no competing interests.

## Authors' contributions

BG and SA conceived of the experiments, SA designed the oligo pool, built the Vex-seq library, and analyzed data. LZ performed cell culture experiments. SA and BG wrote the manuscript. All authors read and approved the final manuscript.

